# Development and evaluation of a loop-mediated isothermal amplification (LAMP) method for detection of the potato wart pathogen *Synchytrium endobioticum*

**DOI:** 10.1101/2022.03.03.482896

**Authors:** Junye Jiang, Will Feindel, David Feindel, Stacey Bajema, Jie Feng

## Abstract

Potato (*Solanum tuberosum*) wart, caused by the biotrophic fungal pathogen *Synchytrium endobioticum*, is a serious disease of cultivated potatoes. A rapid and accurate method for wart diagnosis is needed. In this study, we described the development and evaluation of a loop-mediated isothermal amplification (LAMP) method for the detection of *S. endobioticum*. Four sets of LAMP primers were designed. Their sensitivity was tested on a gBlock and the specificity of the most sensitive primer set (LAMP1) was verified by *in silico* analysis and on DNA of non-target species. The LAMP1 primer set showed a detection limit at 7-8 gBlock DNA molecules per reaction. On sensitivity, the LAMP was comparable to qPCR (SYBR green- and probe-based) and about 10 times more sensitive than conventional PCR. Considering the convenience of operation, the LAMP protocol is accurate and robust and should be recommended for the routine lab testing for potato wart.

## Introduction

Potato (*Solanum tuberosum*) is the fourth most important crop in the world after wheat (*Triticum aestivum*), rice (*Oryza sativa*) and maize (*Zea mays*) (FAO, 2018). In Canada, a new record of potato production was reported with 6.9 million tons in 2021, rising up 18.2% from 2020 (Statistics Canada, 2021). In 2019/2020, Canada exported potatoes with a value of 1.93 billion CAD (Potato Market Information Review: 2019-2020). These statistics address the importance of potato as an edible crop in Canada and worldwide.

The wart, described as the potato cancer by researchers, is a disease caused by an obligate parasite *Synchytrium endobioticum*. Once infected by this pathogen, potato plant shows symptoms on all parts except for the root. On the tuber or stolon, the eyes can develop cauliflower-like swellings, which highly affect the potato quality (Franc 2007). *Synchytrium endobioticum* does not develop mycelium during its lifecycle. The resting sporangia can survive in soil up to 50 cm in depth and for 40 to 50 years without germination (Franc 2007). The disease is devastating to the potato industry, for example, wart disease documented in Prince Edward Island, Canada in 2000 resulted in 30 million CAD just for one year (Clark et al. 2007).

Timely and accurate detection of *S. endobioticum* prior to wide spread of the disease is important for an effective management. In the past 20 years, polymerase chain reaction (PCR)-based methods for potato wart diagnosis were developed and applied in lab tests. A set of conventional PCR primers that can specifically amplify a 472-base pair (bp) DNA fragment of *S. endobioticum* were developed by van den Boogert et al. (2007), which was followed by the development of two sets of more specific and sensitive probe-based quantitative PCR (qPCR) primers (van Gent-Pelzer et al. 2010; Smith et al. 2014). These advances in technology facilitated the rapid and proper diagnosis of potato wart. However, the PCR-based methods are costly and time consuming. Therefore, an alternative and more convenient diagnostic method is needed. The loop-mediated isothermal amplification (LAMP) is a simple technique which takes the advantage of four or six primers and requires only a single step of incubation (Notomi et al. 2000). Compared to PCR-based techniques, LAMP has its advantage in specificity due to the detection of six regions on DNA sequences by the primers, as well as the higher sensitivity (Notomi et al. 2000). In addition, conducting a LAMP experiment does not need specialized equipment such as a thermocycler required by PCR experiments. The results of LAMP can be visually detected and therefore LAMP can be applied with portable equipment.

LAMP diagnostic protocols have been developed for different plant diseases (Huang et al. 2017; Sarkes et al. 2020; Yang et al. 2021; Sun et al. 2022). However, it has not been applied on the detection of *S. endobioticum*. The objective of this study was to develop a LAMP protocol for *S. endobioticum* detection and compare its efficiency with PCR and qPCR. In Alberta, Canada, no potato wart has been detected. Thus our protocol is in the process of verification on positive plant samples by collaborating groups in other provinces. Nevertheless, results from this part of study will be the foundation for the LAMP testing of potato wart in Alberta.

## Materials and methods

### Chemicals and standard techniques

All chemicals and instrument were purchased from Fisher Scientific Canada (Ottawa, ON) unless otherwise specified. All primers, probes and gBlocks were synthesized by Integrated DNA Technologies Canada (Toronto, ON). All DNA extraction was conducted using the DNeasy Plant Pro kits (Qiagen Canada, Toronto, ON). Conventional PCR and LAMP were performed in a ProFlex PCR system. qPCR was performed in a CFX96 touch real-time PCR detection system (Bio-Rad Canada, Mississauga, ON).

### Design of LAMP, PCR and qPCR primers and gBlock

The ribosome DNA sequence (GenBank: KF160871) of a Canadian *S. endobioticum* strain was retrieved from the National Center for Biotechnology Information (NCBI) database (https://www.ncbi.nlm.nih.gov). A 479-bp fragment at the 3’ end of KF160871 was selected. Using this sequence as a query, Blastn was conducted against the nucleotide collection (nr/nt) and whole-genome shotgun contigs (wgs) database, which indicated the availability of the counterpart rDNA sequences in eight *S. endobioticum* strains (confirmed on March 3, 2022). Alignment of the eight sequences indicated that they shared 472 bp (the smallest partial sequence is 472 bp). The sequences of this 472 bp from the eight strains were identical, except for one nucleotide (nt) replacement in one strain. Based on this 472-bp sequence, four sets of LAMP primers (LAMP1 to LAMP4) were designed using an online tool developed by New England Biolabs (https://lamp.neb.com). To compare the efficiency of LAMP and PCR, a pair of primers (qcF/qcR) was designed on this 472-bp sequence by the Primer Blast software (https://www.ncbi.nlm.nih.gov/tools/primer-blast), which was used in conventional PCR and SYBR green-based qPCR. Furthermore, a set of published primers/probe, namely ITS2F/ITS2R/Probe (van Gent-Pelzer et al. 2010), also located within the 472-bp sequence, was used in probe-based qPCR. The 479-bp fragment on KF160871, embracing the 472-bp sequences and all the above-mentioned primers, was synthesized as a gBlock. The locations of all primers were shown in Fig. 1 and all primer sequences were listed in Table 1.

**Fig. 1.**
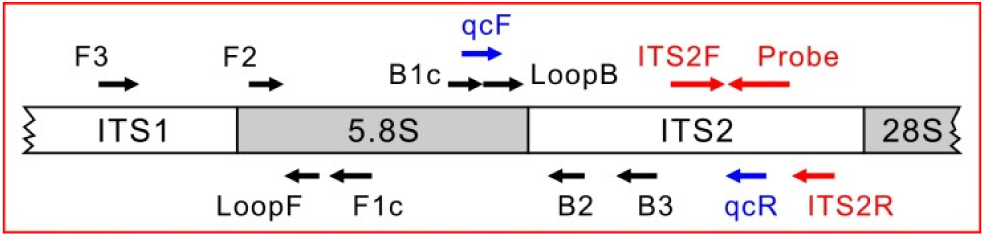
The gBlock fragment of the ribosome DNA region of *Synchytrium endobioticum*. All primers used in this study were indicated. Arrows indicate the 5’ to 3’ direction of the sequences.

**Table 1.**
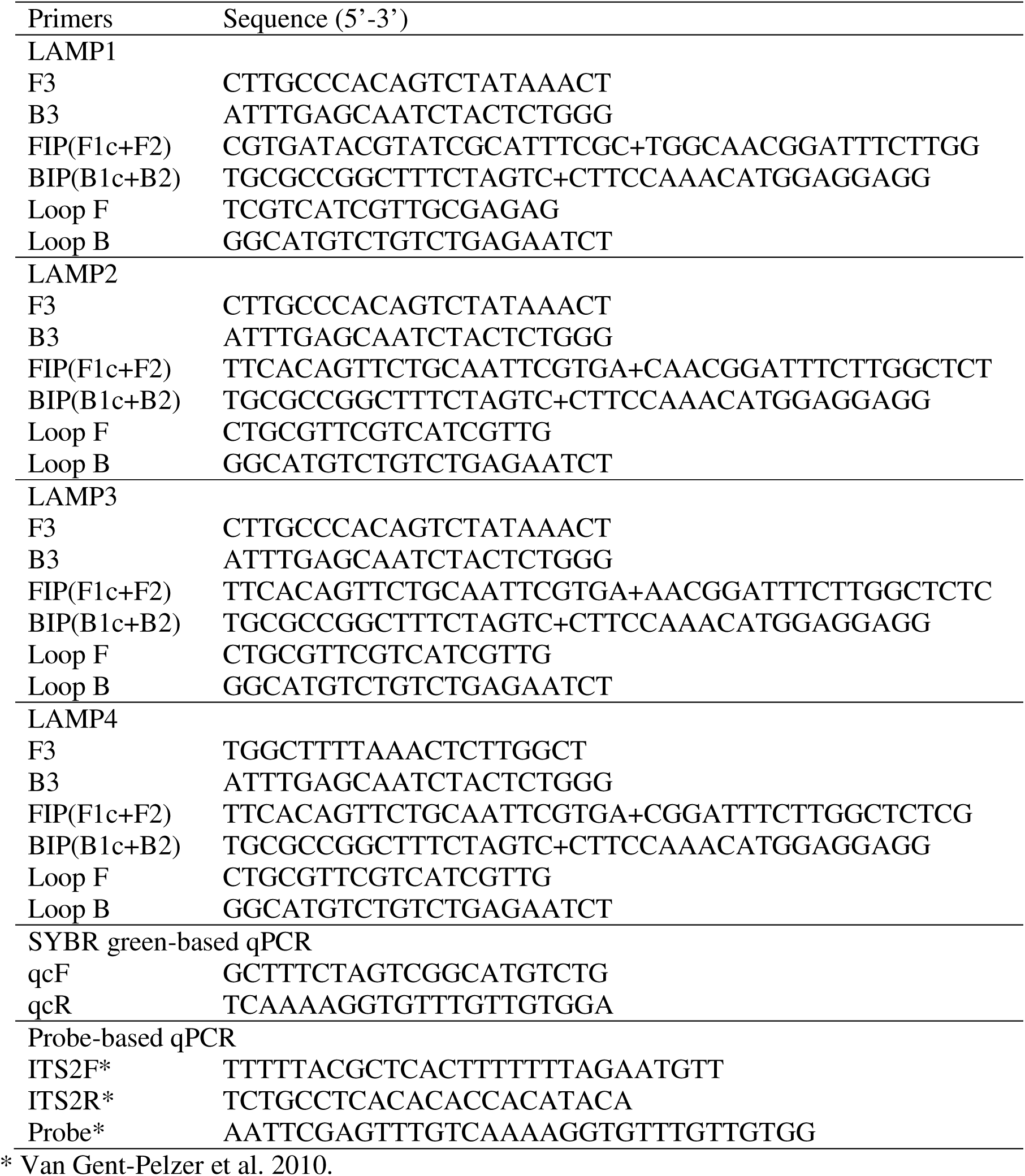
Primers used in this study

### LAMP reaction

All LAMP reactions were conducted in the WarmStart colorimetric LAMP master mix (NEB Canada, Whitby, ON). Each reaction was 25 µL in total volume containing 1 µL template. Quantities of primers in each reaction followed the NEB’s instructions for the master mix. All reactions were conducted in individual 200-µL PCR tubes. The reaction program consisted of only one step in which the tubes were incubated at 60^°^C for 50 minutes. After the incubation, the reactions were checked visually and the results were recorded by photography.

### PCR and qPCR reactions

Each reaction of PCR and qPCR was 25 µL in total volume containing 1 µL template and 0.25 µM of each primer and 0.125 µM of probe (probe-based qPCR only). PCR was conducted in Promega PCR master mix (Promega, Madison, WI). SYBR green-based qPCR and probe-based qPCR was conducted in SsoAdvanced universal SYBR green supermix (Bio-Rad Canada) and SsoAdvanced universal probe supermix (Bio-Rad Canada), respectively. The PCR program consisted of an initial denaturation at 95^°^C for 5 min, followed by 40 cycles of denaturation at 95^°^C for 30 s, annealing at 58^°^C for 45 s and extension at 72^°^C for 45 s, and a final extension at 72^°^C for 10 min. The qPCR program consisted of an initial denaturation step of 95^°^C for 2 min, followed by 40 cycles of 5 s at 95^°^C and 1 min at 60^°^C.

### Sensitivity test on gBlock

From the stock solution of the gBlock, a set of 10× serial dilutions were prepared from 6 × 10^7^ to 0.6 molecules/µL (nine solutions). Using the serial dilutions as templates, LAMP and PCR were conducted to evaluate the sensitivity of the primers/probe.

### Specificity test of LAMP

The specificity of F3/B3 and F2/B2 in the LAMP1 primer set was tested by primer blast (https://www.ncbi.nlm.nih.gov/tools/primer-blast/index.cgi). In addition, genomic DNA from potato (cv. Russet Burbank) tubers, *Plasmodiophora brassicae* and fungi other than *S. endobioticum* was used as templates. The fungi were 23 species isolated from fields in Alberta and maintained in our culture collection, including *Alternaria solani, Botrytis cinerea, Colletotrichum coccodes, Fusarium avenaceum, F. chlamydosporum, F. culmorum, F. equiseti, F. graminearum, F. incarnatum, F. lateritium, F. oxysporum, F. poae, F. sambucinum, F. solani, Leptosphaeria biglobosa, Neonectria candida, Pyrenophora teres, Pythium intermedium, Rhizoctonia solani, Sclerotinia sclerotiorum, Verticillium albo-atrum, V. longisporum* and *V. nonalfalfae*. All fungal species were cultured on 4% PDA plates and mycelia were collected. In addition, soil samples were collected from a potato field at the Crop Diversification Center North, Edmonton, AB. One gram of the soil sample was cultured in 30-mL of YPG media (1% yeast extract, 1% peptone and 2% glucose) and 2% LB broth. The cultures were shaken at 200 rpm at 25^°^C (YPG) or 37^°^C (LB) overnight and the pellets from the cultures were collected by centrifugation. DNA was extracted from the pellets, as well as mycelia, potato tubers and a clubroot gall on canola (for *P. brassicae*). Twenty ng of the resultant DNA was used in each LAMP reaction.

### Statistics

The lowest number of gBlock molecules in a positive reaction was calculated according to Poisson distribution by online calculator (https://stattrek.com/online-calculator/poisson.aspx). In each qPCR reaction, the mean of the quantification cycle (Ct) values from three technical repeats was calculated and treated as one data point. The qPCR standard curves were constructed by regression analysis using the R software (https://cran.r-project.org).

## Results

### Sensitivity test

Among the four sets of LAMP primers, the LAMP1 primer set showed the highest sensitivity, with the low limit of positive detection at 60 gBlock molecules per reaction, which was at least 10 times lower than the other three sets of primers (Fig. 2). SYBR green-based qPCR using the primer pair qcF/qcR and the probe-based qPCR using ITS2F/ITS2R/Probe of van Gent-Pelzer et al. (2010) resulted in a low limit of positive detection at 6 gBlock molecules per reaction, with a Ct value of 35.8 and 35.6, respectively (Fig. 2). The primer pair qcF/qcR was also used in a conventional PCR against the gBlock serial dilutions, from which the low limit to amplify a product (as a visible band on agarose gel) was 60 molecules per reaction (Fig. 3).

**Fig. 2.**
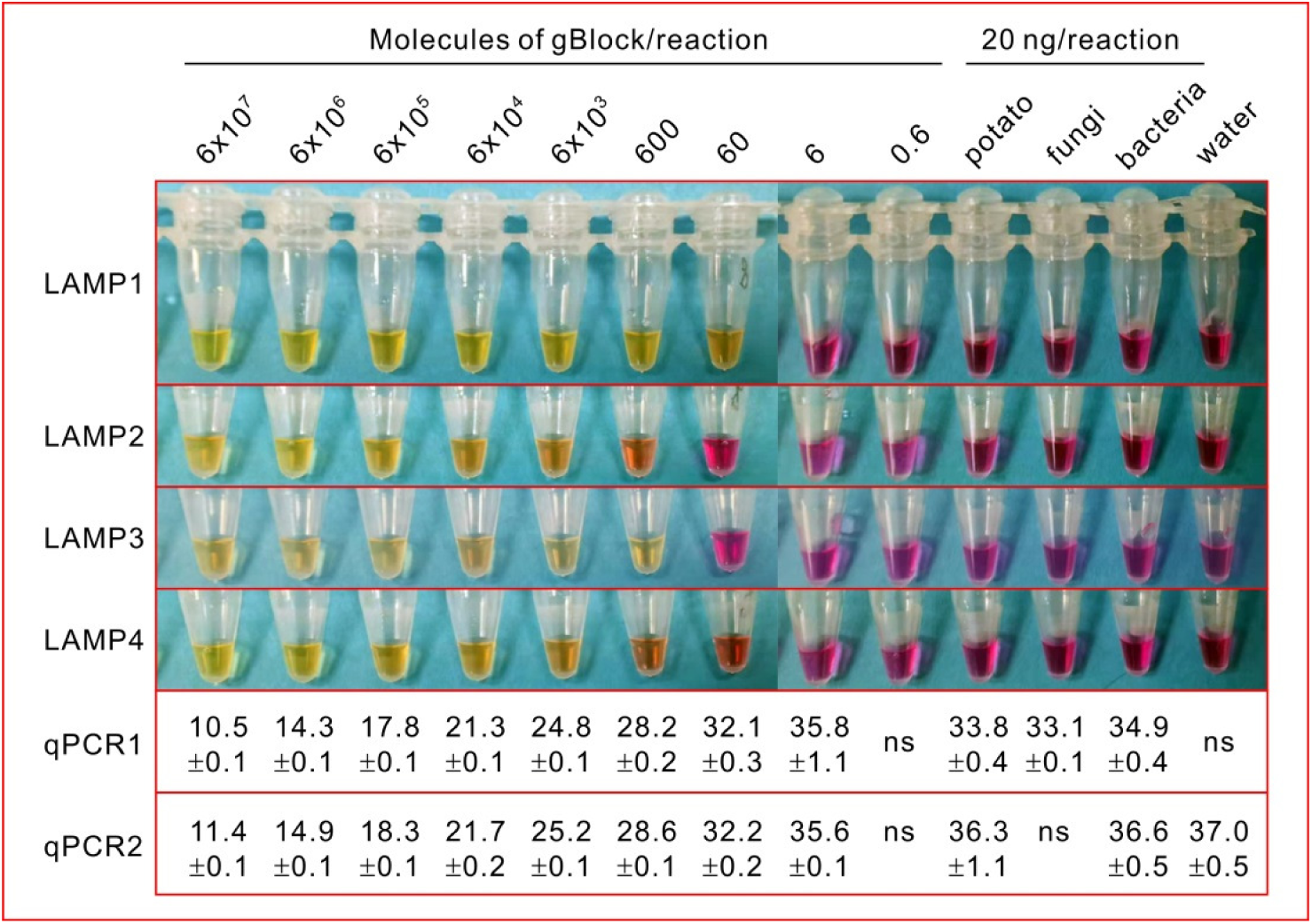
Sensitivity test of four sets of LAMP primers and two sets of qPCR primers on the gBlock. qPCR1 (primers qcF/qcR) and qPCR2 (primers ITS2F/ITS2R and Probe) were designed by this study and by van Gent-Pelzer et al. (2010), respectively. In LAMP assays, yellow or light orange color indicates a positive reaction and pink or dark orange color indicates a negative reaction. In qPCR assays, the quantification cycle (Cq) values are shown as mean ± standard deviation (n=3). ns, no signal.

**Fig. 3.**
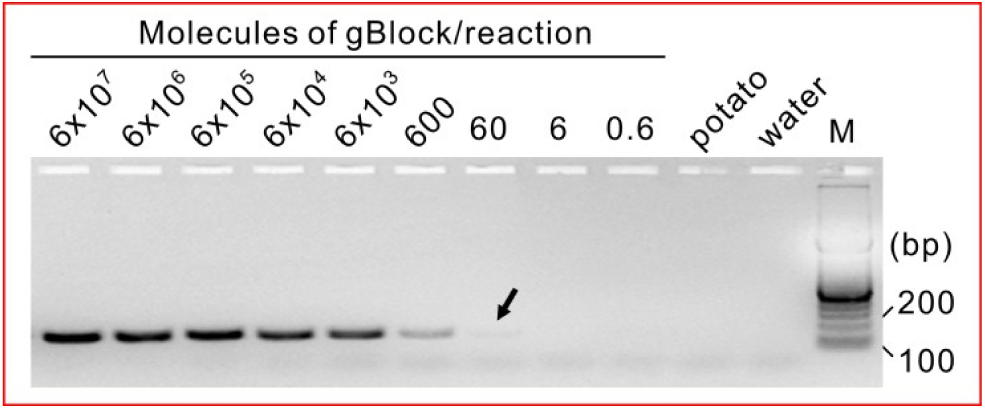
Sensitivity test of conventional PCR using primers qcF/qcR on the gBlock. The arrow indicates a weak band. M, DNA ladder.

### Estimation of the lowest target molecule number per reaction in a positive LAMP reaction

The sensitivity of the LAMP1 primers was further investigated by repeated running LAMP reactions of 60, 6 and 0.6 gBlock molecules per reaction. All the eight repeats of 60 molecules per reaction (Fig. 4a) and none of the eight repeats of 0.6 molecules per reaction (Fig. 4b) produced positive signals. For the 37 repeats of 6 molecules per reaction, in addition to yellow and pink colors, orange color was observed from some reactions (Fig. 4c). If we defined the light orange as positive and dark orange as negative reaction, among the 37 repeats, 13 were positive and 24 were negative (Fig. 4). Thus the proportion of positive reactions was 13/37 = 35.1%. According to the Poisson distribution, the probabilities that a 1-µL aliquot of a 6 molecules/µL solution contains ≥7 and ≥8 molecules are 39.4% and 25.6%, respectively. Therefore, the lowest molecule number for a positive reaction by LAMP1 was estimated as 7 or 8, which was more sensitive than conventional PCR.

**Fig. 4.**
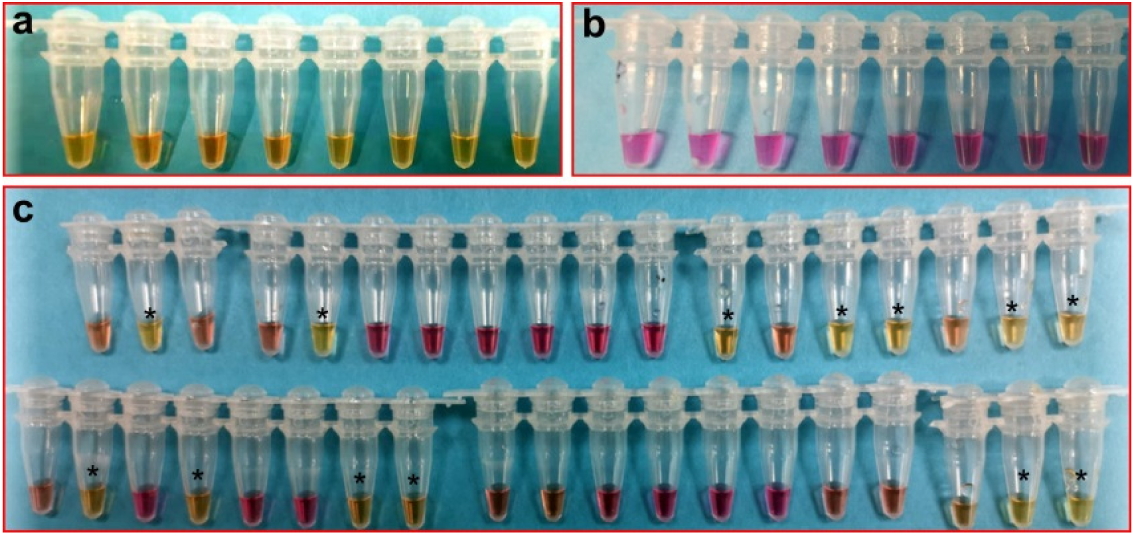
Evaluation the low limit of the LAMP for a positive detection on the gBlock. a, eight repeats of 60 molecules/reaction; b, eight repeats of 0.6 molecules/reaction; c, 37 repeats of 6 molecules/reaction. Yellow or light orange color indicates a positive reaction (marked by a *); pink or dark orange color indicates a negative reaction.

### Specificity test

Primer blast search against the NCBI nr databases indicated that the sequences of the primer pairs F3/B3 and F2/B2 of the LAMP1 primer set were exclusively present in the *S. endobioticum* rDNA region. DNA from potato, the clubroot pathogen *P. brassicae*, 23 fungal species and microorganisms present in the soil of potato field were used in the specificity test of the LAMP1 primer set. All reactions remained pink (data not shown). These results indicated that LAMP1 is specific to *S. endobioticum*.

## Discussion

In plant disease diagnostics, sensitivity and specificity are the two most important parameters to be tested for a protocol. With a relatively high sensitivity and specificity, the qPCR technique has become a popular strategy in diagnostics. However, the high expense of equipment and the requirement of long-time training for employees restricted the qPCR usage in many labs. Therefore, researchers have been seeking alternative techniques affordable and convenient to operate in most labs. The LAMP (Notomi et al. 2000) technique provides a useful method to guarantee both sensitivity and specificity, and more convenient and cheaper than qPCR.

One important advantage of using gBlock as a template is that the molecule number needed in a positive reaction can be estimated, especially for an obligate parasite such as *S. endobioticum* which cannot be easily quantified in a DNA mixture with its host. In the sensitivity test, at the gBlock concentration of 6 molecules/µL, 13 of 37 repeats showed positive reactions, accounting for 35.1%. The numbers of DNA particles in different aliquots could be described as a Poisson distribution (Wang and Spadoro 1998), according to which, the lowest molecule number to be detected by the LAMP1 primers was 7 or 8 per reaction. In the qPCR assays, the results should be considered as technically negative when the Ct value is larger than 36 (Okiro et al. 2019). In the current study, the gBlock molecule number per reaction was 6 when Ct value was smaller than 36 in qPCR experiments. Therefore, the LAMP sensitivity is comparable with both probe-based and SYBR green-based qPCR assays. Similar sensitivities between LAMP and qPCR have been reported in previous studies (Okiro et al. 2019; Sarkes et al. 2020). In the conventional PCR test, a very weak band was observed at the molecule number of 60 per reaction, which indicated the LAMP was about 10 times more sensitive than the conventional PCR. Similarly, the conventional PCR was reported to be less sensitive than LAMP assay in diagnosis of other diseases (Khan et al. 2018; Sarkes et al. 2020; Arizala et al. 2022). In addition, the LAMP assay eliminates the down-stream work needed in conventional PCR assay such as gel electrophoresis. All the information described proves that the LAMP1 primer set showed great sensitivity in diagnosis for potato wart disease.

The LAMP primers were designed at an rDNA region where specific qPCR primers were located (van Gent-Pelzer et al. 2010; Smith et al. 2014). With the rapid development of sequencing technology, a sufficient amount of genomic information became available, which provides a useful tool and has proven to be an essential component for the development of a robust and reliable diagnostic assay (Manjunatha et al. 2018; Karim et al. 2019; Arizala et al. 2022). In this study, the *in silico* specificity test showed that the LAMP primers are exclusively present in the *S. endobioticum* rDNA region, suggesting they are unique and highly conserved in the natural evolution process. The *in vitro* specificity test of LAMP assays did not show any positive reaction on all the non-target organisms, confirming the high specificity of the LAMP primers.

It has not escaped our notice that the SYBR green-based qPCR had a compromised specificity, as shown in Fig. 2 (qPCR1), Ct values smaller than 36 were produced from non-target genomic DNA. This is likely due to the fact that only two specific regions carried by the two primers were present in the SYBR green-based qPCR. In contrast, LAMP has six specific regions carried by the four primers, which guarantees LAMP a general higher specificity compared to PCR-based techniques.

Another advantage of LAMP assay is time and cost saving. In the current study, one experimental ran only for 50 minutes and all the assays were conducted as one step in a PCR thermocycler. For labs without a PCR thermocycler, the LAMP assay could be conducted in a heat block or a water bath, which will greatly reduce the financial expense and even could be applied in the field by the farmers directly.

In conclusion, we developed and evaluated a LAMP assay for the rapid and accurate detection of the potato wart pathogen *S. endobioticum*. The lowest molecule number for a positive detection in one reaction was 7 or 8, which was highly sensitive. The specificity test indicated a unique and conserved character of this diagnostic tool. We do believe this LAMP protocol will be beneficial to the proactive diagnosis and management of potato wart disease.

## Funding

Financial support was received from Results Driven Agriculture Research (RDAR) for a project entitled establishment of a platform for rapid diagnosis of current and potential potato diseases in Alberta.

## Notes

### Competing Interest Statement

The authors have declared no competing interest.

